# The absence of the autoimmune regulator gene (*AIRE*) impairs the three-dimensional structure of medullary thymic epithelial cell spheroids

**DOI:** 10.1101/2021.08.11.455994

**Authors:** Ana Carolina Monteleone-Cassiano, Janaina A. Dernowsek, Romario S. Mascarenhas, Amanda Freire Assis, Dimitrius Pitol, João Paulo Mardegan Issa, Eduardo A. Donadi, Geraldo Aleixo Passos

## Abstract

Besides controlling the expression of peripheral tissue antigens, the autoimmune regulator (*AIRE*) gene also regulates the expression of adhesion genes in medullary thymic epithelial cells (mTECs), an essential process for mTEC-thymocyte interaction for triggering the negative selection in the thymus. For these processes to occur, it is necessary that the medulla compartment forms an adequate three-dimensional (3D) architecture, preserving the thymic medulla. Previous studies have shown that *AIRE* knockout (KO) mice have a small and disorganized thymic medulla; however, whether *Aire* influences the mTEC-mTEC interaction in the maintenance of the 3D structure has been little explored. Considering that AIRE controls cell adhesion genes, we hypothesized that this gene affects 3D mTEC-mTEC interaction. To test this, we constructed an *in vitro* model system for mTEC spheroid formation, in which cells adhere to each other, establishing a 3D structure. The effect of *Aire* on mTEC-mTEC adhesion was evaluated by comparing *AIRE* wild type (*AIRE*^*WT*^) versus Aire KO (*AIRE*^*-/-*^) mTECs. Considering the 3D spheroid model evaluated, we reported that the absence of *AIRE* disorganizes the 3D structure of mTEC spheroids, promotes a differential regulation of mTEC classical surface markers, and modulates genes encoding adhesion and other molecules.

## Introduction

The three-dimensional (3D) structure of the thymus is composed of two histologically distinct but closely related compartments. The cortex is mainly formed by cortical thymic epithelial cells (cTECs), and the medulla is primarily formed by medullary thymic epithelial cells (mTECs), which in mice present the CD45^-^, EpCam^+^, Ly51^+^, UEA1^+^ phenotype [1-3]. Other cell types include thymic seeding progenitors, thymocytes at different stages of maturation, including double-negative (CD4^-^CD8^-^), double-positive (CD4^+^CD8^+^), and single-positive (CD4^+^CD8^-^ and CD4^-^CD8^+^) thymocytes, thymic B cells, macrophages, dendritic cells, and Hassall’s corpuscles [2].

The thymic 3D architecture offers a particular microenvironment for the positive and negative selection of developing thymocytes [4-6]. The thymic cortex is implicated in the positive selection of thymocytes that express functional T-cell receptors (α/β TCRs). The medulla is involved in the negative selection of autoreactive thymocytes, which avidly recognize self-peripheral tissue antigens (PTAs) presented on the surface of mTECs. This unique pattern of gene expression allows mTECs to express more than 19,000 protein-encoding genes, including the “ectopic” genes that encode PTAs, a phenomenon known as promiscuous gene expression (PGE) [4, 7-10]. At present, no other cell type is known that expresses such a large set of genes [11-13]. The positive and negative selection processes are crucial for the induction of central immune tolerance that prevents aggressive autoimmunity [9, 14].

The autoimmune regulator gene (AIRE) is the primary transcriptional regulator of PTAs in mTECs, whose encoded Aire protein unleashes stalled RNA Pol II in the chromatin to proceed with the elongation phase of transcription [15, 9, 10]. A second controller of PGE in mTECs is the forebrain embryonic zinc finger-like protein 2 (*FEZF2*) that plays a role as a classical transcription factor and thus binds directly to DNA in specific promoter regions [4,16].

Given the non-specificity of RNA Pol II in transcribing genes and that *AIRE* protein in mTECs unleashes this enzyme on chromatin, AIRE ends up controlling the expression of a diverse range of mRNAs. In addition to PTAs, AIRE also regulates the expression of genes involved in cell adhesion, such as the extracellular matrix (ECM) constituent Lama1, the CAM family adhesion molecules Vcam1 and Icam4, which control mTEC-thymocyte adhesion [17, 18]. Cell adhesion corresponds to an essential biological process in the structure and function of the thymus. The adequate adhesion of TECs increases the efficiency for T-cells to develop [19, 20]. Besides adhesion molecules, the formation of a dense cellular network is necessary for the maturation of thymocytes, which is composed of ECM proteins such as laminins, integrins, collagens, and fibronectins, as well as soluble molecules such as hormones, cytokines, chemokines, and growth factors that are also mediated by *AIRE* [21].

*AIRE*^*-/-*^ mice have small and disorganized medulla of the thymus [22], and little is known about whether Aire influences the mTEC-mTEC adhesion in a 3D thymus structure. Two experimental strategies enable the study of the *in vitro* formation of the thymus’s 3D structure that reproduces its microenvironment with the extracellular matrix and thymic epithelial cells. One of these strategies is the thymus re-aggregation [19, 20], and the other is the 3D organotypic culture [23]. In the re-aggregation model, the thymus tissue is devoid of cells (decellularized). Still, it retains most of the microenvironment of the extracellular matrix that could support the re-aggregation between TECs. When transplanted into athymic nude mice, the re-aggregate thymus organoids can receive lymphocyte progenitors derived from bone marrow and develop a diverse and functional T-cell repertoire [19, 20].

In the search for an experimental model that could mimic the thymic microenvironment closest to that found *in vivo*, Pinto et al. [23] adapted a 3D organotypic co-culture, preserving the main characteristics of mTECs, such as proliferation and differentiation. This strategy helped to identify molecular components and pathways involved in the mTEC differentiation and promiscuous gene expression. Other studies used stem cells to form “thymospheres,” permitting the study of the factors necessary for thymocyte development [24, 25]. Recently, a 3D culture substrate has been reported that allowed TECs to survive and proliferate, using electrospun fibrous meshes (eFMs) functionalized with fibronectin. The mTECs presented increased proliferation, viability, and protein synthesis when cultured on fibronectin-functionalized eFMs (FN-eFMs) [26].

In the present study, we asked whether *AIRE* regulates the adhesion between mTECs during the in *vitro* formation of a 3D structure. For this, we developed a model system in which mTECs are grown in non-adherent agarose micro-wells and adhere to form spheroids. The comparison between *AIRE* wild type (*AIRE*^*WT*^) versus *AIRE*^*-/-*^ mTECs allowed us to evaluate the influence of *AIRE* in the adhesion between these cells during the spheroid formation.

## Materials and methods

### Medullary thymic epithelial cell lines

We use the *AIRE*^*WT*^ murine (*Mus musculus*) mTEC 3.10 line (EpCAM^+^, CD45^-^, Ly51^−^, UEA-1^+^), as previously described [17, 27, 28], and the *AIRE*^*-/-*^ mTEC 3.10E6 cell clone. The mTEC 3.10E6 clone obtained by the CRISPR-Cas9 system [18] was characterized as a carrier of indel mutations affecting both *AIRE* alleles (compound heterozygosis). In the *AIRE* allele 1, there were two types of mutations: a T>G substitution (mRNA nucleotide, nt, position 351) followed by a nine-bp deletion (GCTGGTCCC, mRNA nt positions 352–360) that transcribed a 1,647 nt *AIRE* mRNA. In allele 2, a single G deletion at mRNA nt position 352 transcribed a 1,655 nt *AIRE* mRNA. As both alleles of the mTEC 3.10E6 clone produced a nonfunctional *AIRE* protein, they were considered knock out; i. e., *AIRE*^*-/-*^.

### Spheroid formation

We use a precast agarose mold with non-adherent 600 µm diameter microwells, making the mTEC cells adhere once seeded in these compartments. The *AIRE*^*WT*^ mTEC 3.10 and the *AIRE*^*-/-*^ mTEC 3.10E6 cell lines were initially cultured as monolayers in RPMI 1640 medium (Gibco, Darmstadt, Germany) supplemented with 10% inactivated fetal bovine serum in 75 cm^2^ polystyrene plastic bottles (Corning, New York, NY) in an incubator at 37° C with 5% CO_2_ atmosphere. After acquiring confluence, mTECS were trypsinized and seeded in agarose molds, using 2% low electroendosmosis agarose (Sigma-Aldrich, Saint Louis, MO), sterilized in 70% ethanol, washed twice in sterile PBS, and irradiated under germicidal UV light for 15 minutes. Spheroid growth was observed through a Cytosmart® (Lonza Group AG, Basel, Switzerland) inverted microscope to produce a real-time movie of the culture.

### Growth curve

To better characterize the 3D culture model, we quantified the cells that form the spheroids at different time points (determinations at every 12 h). At each time point, the spheroids were removed from the agarose microwells and dissociated using trypsin. Isolated cells were counted using a Cellometer® Auto T4 Bright Field Cell Counter (Nexcelom Bioscience, Lawrence, MA). Triplicates were performed for each time point. We have thus drawn a spheroid growth curve to identify the exponential, stationary growth, and decline phases.

### Histological analysis of spheroids

The spheroids were fixed in 10% formaldehyde buffered in PBS, then dehydrated, passing through a battery of aqueous ethanol solution (75 to 100% ethanol) and included in historesin (HistoResin standard kit, Biosystems Switzerland AG, Muttenz, Switzerland). The resin blocks were cut to 1 µm thickness and deposited on microscope slides. After deparaffinization and rehydration, the slides were stained with hematoxylin-eosin (H&E) for further microscopic examination of the spheroid morphology.

### Live-dead assay

We used the LIVE/DEAD® Viability/Cytotoxicity Kit (Thermo-Fisher, Waltham, MA) to assess the proportion of live and dead cells in the spheroids, following the manufacturer’s instructions.

### Scanning electron microscopy (SEM)

Spheroids were fixed with 1.6% glutaraldehyde in 0.2 M buffered (pH 7.4) sodium cacodylate overnight at 4 °C, then washed three times with 0.2 M sodium cacodylate buffer (pH 7.4). Spheroids were immersed into 1% osmium tetroxide (Sigma-Aldrich) for secondary fixation in 0.2 M buffered (pH 7.4) sodium cacodylate. Fixed spheroids were subsequently dehydrated in 25-100% ethanol and immersed in hexamethyldisilazane (HMDS, Sigma-Aldrich) for 10 minutes at room temperature. The spheroids were freeze-dried until the HMDS had evaporated, mounted, coated with palladium gold, and examined under a scanning electron microscope (Jeol JSM -6610 LV, Hitachi High-Tech America, Tokyo, Japan) at an acceleration voltage of 5 kV. The SEM procedures were done at the Electron Microscopy Facility, Ribeirão Preto Medical School, University of São Paulo, Ribeirão Preto, SP, Brazil.

### Flow cytometry

We used a FACSCanto™ II (Becton Dickinson, Franklin Lakes, NJ) flow cytometer to evaluate the medullary phenotype of *AIRE*^*WT*^or *AIRE*^*-/-*^ spheroids. A sample of 1 × 10^6^ mTECs separated from spheroids were labeled with 1:250 dilution of anti-mouse CD45-PerCP, anti-mouse CD326 (EpCam), anti-mouse Ly51-PE (BD Biosciences, San Jose, CA), anti-mouse CD80-APC, and anti-mouse MHCII-PE antibodies in a final volume of 200 μL of cell suspension. Labeling was also performed with a 1:250 dilution of Lectin agglutinin I (Ulex europaeus UEA-I)-FITC (Vector Labs, Burlingame, CA).

### Total RNA preparation and cDNA synthesis

Total RNA of *AIRE*^*WT*^ or *AIRE*^*-/-*^ spheroids was prepared using the mirVana kit® (Ambion, Austin, TX) according to the manufacturer’s instructions. Microfluidic electrophoresis evaluated RNA integrity using Agilent RNA 6000 nanochips and an Agilent 2100 Bioanalyzer (Agilent Technologies Santa Clara, CA). Only RNA samples that were free of proteins and phenol and had an RNA Integrity Number (RIN) ≥7.0 were selected for cDNA synthesis or RNA sequencing. For cDNA synthesis, the SuperScript® reverse transcriptase enzyme (Invitrogen Corporation, Carlsbad, CA) was used according to the manufacturer’s instructions.

### Transcriptome analysis through RNA-Seq

We followed a protocol previously described by St-Pierre et al. [11]. Briefly, paired-end (2 × 150 bp) sequencing was performed by an Illumina HiSeq 2500 sequencer (Illumina, San Diego, CA), using a TruSeq Stranded Total RNA Library Prep Kit (Illumina). The quality of raw FASTQ sequences was first analyzed through a FASTQC program (https://www.bioinformatics.babraham.ac.uk/projects/fastqc/). Then, FASTQ sequences were mapped to the *Mus musculus* reference genome (mm21) using the STAR 2.5 Spliced Aligner program (https://github.com/alexdobin/STAR), which outputs BAM file containing the sequences and their genomic references and a GTF file with gene annotations used for further determinations of the number of reads per transcript through the HTSeq Count program (http://htseq.readthedocs.io). For each RNA sample analyzed, we recovered a list of genes and their respective number of transcripts that served as input for determinations of the differentially expressed (DE) mRNAs through the EdgeR package (https://bioconductor.org/packages/release/bioc/html/edgeR.html) within the R platform (https://www.r-project.org). EdgeR calculates the fold change (FC) for each mRNA, considering a contrast matrix for a given experimental condition. This study defined WT as a contrast variable and as DE the mRNAs exhibiting a p-value < 0.05 and a false discovery rate (FDR Benjamini–Hochberg correction) FC ≥ 2.0. The DE mRNAs were hierarchically clustered, and a heat map was constructed to evaluate the expression profiling.

### Functional enrichment of differentially expressed mRNAs

The list of the DE mRNAs was analyzed in terms of functional enrichment through the Database for Annotation, Visualization, and Integrated Discovery (DAVID) annotation tool (https://david.ncifcrf.gov/) to the identification of the main biological processes and pathways represented by DE mRNAs. A functional category was considered significant if it comprised at least five mRNAs and a score of p < 0.005 with Benjamini–Hochberg correction.

### Reverse transcription quantitative real-time PCR (RT-qPCR)

The validation of the transcriptional expression of selected DE mRNAs was assayed by reverse transcription-quantitative real-time PCR (RT-qPCR) of the respective cDNAs. The expression level of each target mRNA was normalized to the housekeeping mRNA Hprt, which is commonly used as a reference. The Primer Blast (https://www.ncbi.nlm.nih.gov/tools/primer-blast/) web tool was used to select pairs of oligonucleotide primers spanning an intron/exon junction with an optimal melting temperature of 60°C. The respective sequences were retrieved from the NCBI GenBank database (https://www.ncbi.nlm.nih.gov/). The forward (F) and reverse (R) primer sequences (presented in the 5⍰–3⍰ orientation) were as follows: Hprt (NM_013556.2) F = 5’ GCCCCAAAATGGTTAAGGTT 3’ R = 5’ CAAGGGCATATCCAACAACA 3’, Plcb2 (NM_177568.2) F = 5’TGGAGTTCCTGGATGTCACG3’ R = 5’GCAGGAAGTGGTTGTCTGGA3’, Parvb (NM_133167.3) F = 5’ TCTTTCTTGGGCAAGTTGGG3’ R = 5’ CCATTGGAGAGTTGATGGCG3’, P2rx7 (NM_011027.4) F = 5’
sGCACGAATTATGGCACCGTC3’ R = 5’TAACAGGCTCTTTCCGCTGG3’, Vegfc (NM_009506.2) F = 5’GCTGATGTCTGTCCTGTACCC3’ R = 5’ACTGTCCCCTGTCCTGGTAT3’, Id1 (NM_001369018.1) F= 5’CCTGAACGGCGAGATCAGTG3’ R= 5’AAGTAAGGAAGGGGGACACC3’.

Gene expression was quantified using a StepOne Real-Time PCR System apparatus (Applied Biosystems, Waltham, MA). The analyses were performed using the cycle threshold (Ct) method, which allows for quantitative analysis of the expression of a factor using the formula 2−ΔΔCt, in which ΔCt = Ct target gene − Ct of the housekeeping gene Hprt, and ΔΔCt = ΔCt sample − ΔCt. Experiments were performed in three independent replicates, and statistical analysis of the data was made through the Mann-Whitney two-sided test with 95% interval.

## Results

### The absence of Aire disorganizes the initial phases of spheroid formation

Figure 1A shows the beginning of spheroid growth (0 h), comparing *AIRE*^*WT*^ mTEC 3.10 vs. *AIRE*^*-/-*^ mTEC 3.10E6. The cultures started from the seeding of mTEC cells (0 h) until the complete 3D spheroid formation (24 h). A compact real-time video of the dynamics of the spheroid formation, recorded from 0 to 24 h, is shown in the supplemental material video 1 (available at www.rge.fmrp.usp.br/passos/esferoides). The results show that the mTEC cell lines (*AIRE*^*WT*^ or *AIRE*^*-/-*^) provide morphologic differences during the development of spheroids. The *AIRE*^*WT*^ spheroids consolidate and are firmly compact at 12 h in culture, whereas the *AIRE*^*-/-*^ spheroids showed a delay in cell-cell adhesion and consolidate after 20 h. This finding indicates that in the absence of *AIRE*, mTEC-mTEC adhesion is impaired.

**Figure 1A.**
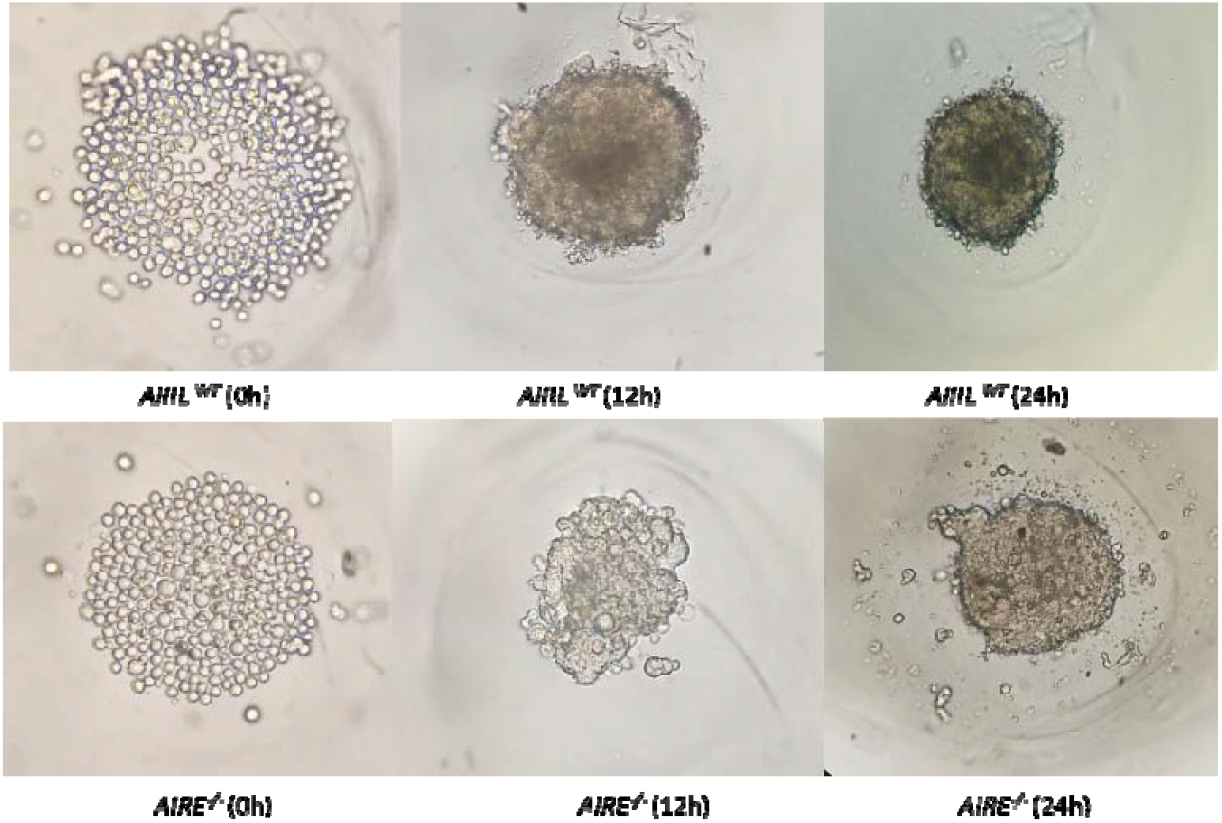

To better characterize the 3D spheroid model system, we constructed a growth curve counting cells at each 12 h of culture. At the first 12 h culture, *AIRE*^*-/-*^ spheroids had a cell count higher than *AIRE*^*WT*^ spheroids. Differently, the development of the *AIRE*^*WT*^ spheroids drew a classical growth curve permitting the identification of the exponential, stationary, and decay phases. The slope of the *AIRE*^*-/-*^ spheroid growth curve indicates that in the absence of Aire, the growth is increased at the beginning, and then the growth abruptly decreases. This suggests that in addition to controlling adhesion between cells, Aire might influence the cell cycle of mTECs (Figure 1B).

**Figure 1B.**
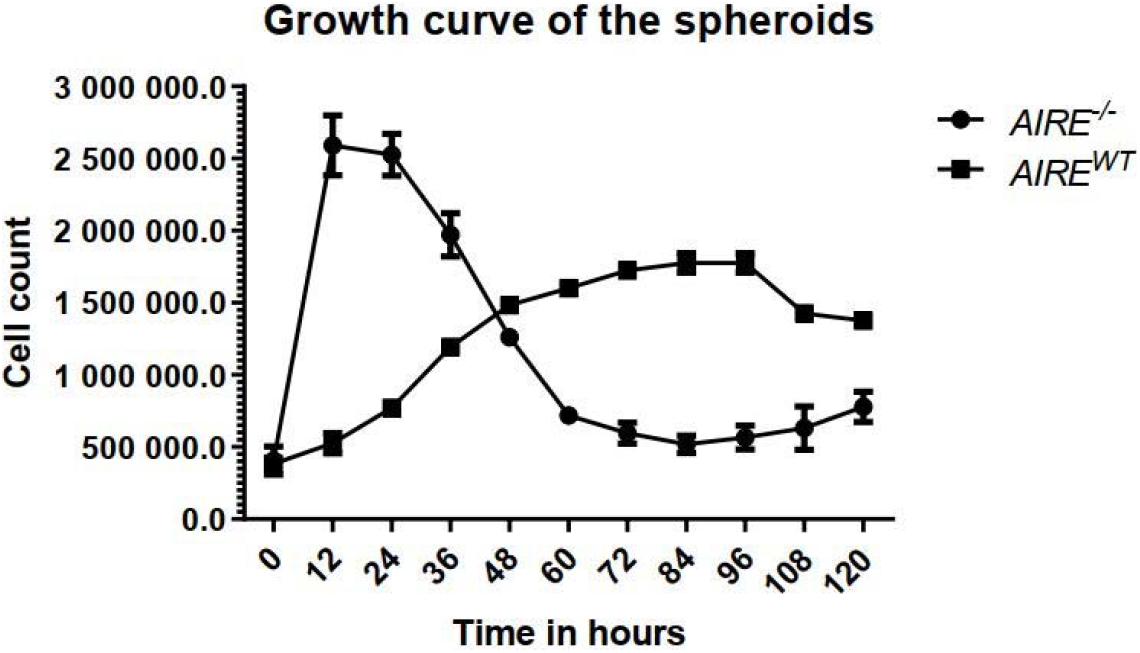

Besides the growth curve, we evaluated the cell viability during spheroid growth. The *AIRE*^*-/-*^ spheroids increased viability during the first 12 h and decreased thereafter, whereas the *AIRE*^*WT*^ progressively reached viability starting from 12 h and maintaining thereafter (Figure 1C).

**Figure 1C.**
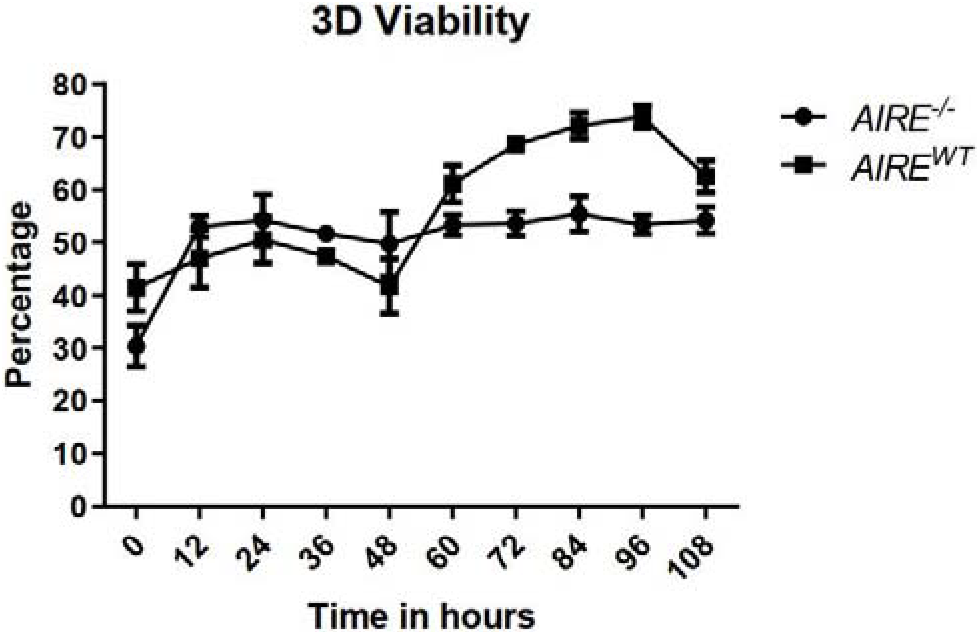

The light microscopy of the 1 µm histologic cuts allowed the observation of the internal structure of spheroids. The *AIRE*^*WT*^ or *AIRE*^*-/-*^ spheroids showed significant differences in their areas, especially at the first 24 h of growth. Although *AIRE*^*-/-*^ spheroids present a larger area than *AIRE*^*WT*^ spheroids. *AIRE*^*-/-*^ cells are dispersed and compact similarly to *AIRE*^*WT*^ spheroids only after 24 h of adhesion (Figure 1D, 1E).

**Figure 1D.**
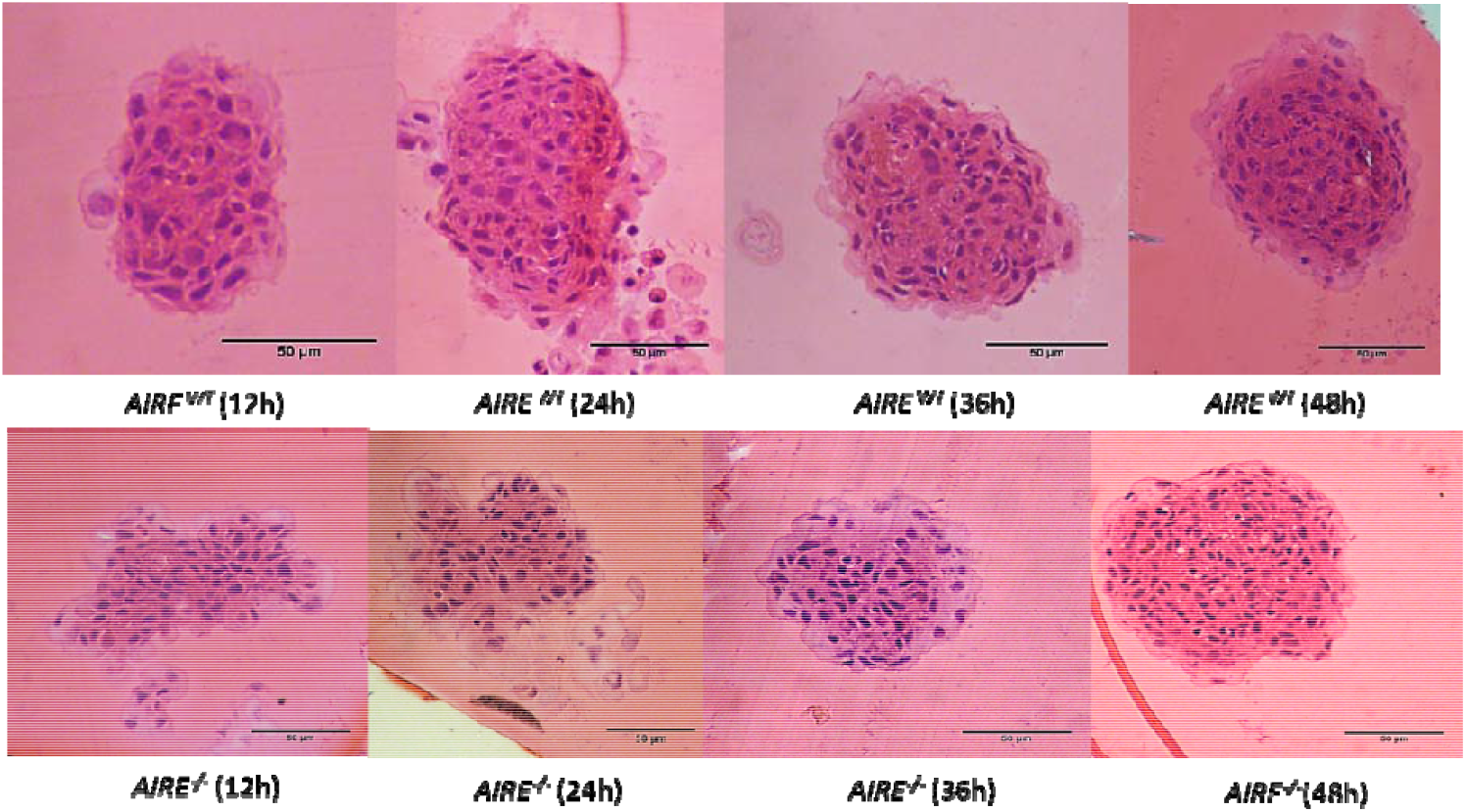

**Figure 1.**
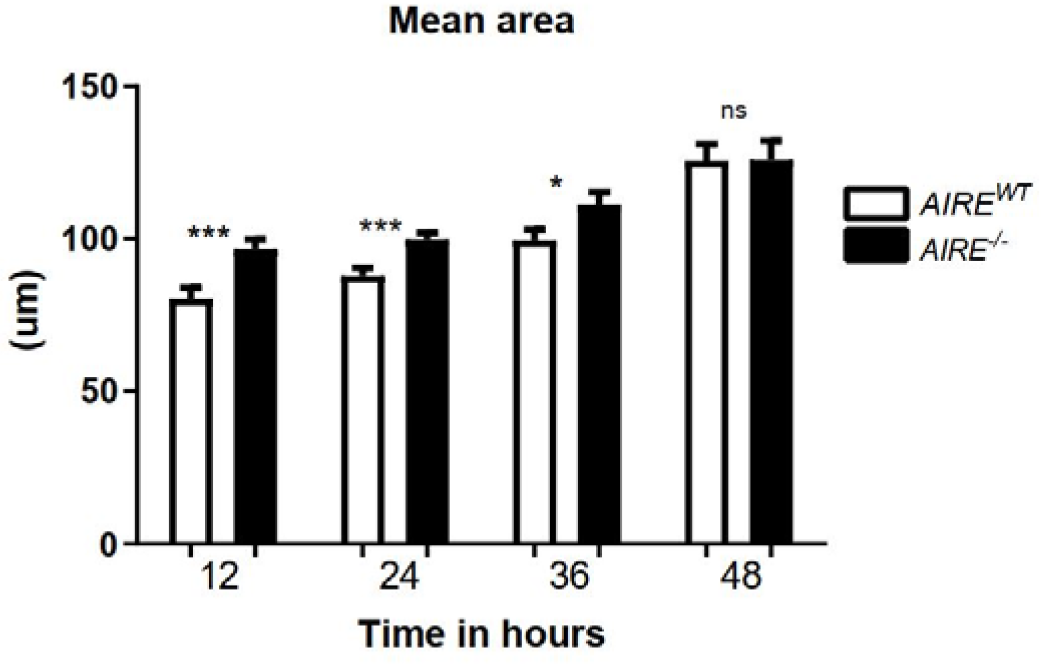
Comparisons between *AIRE*^*WT*^ and *AIRE*^*-/-*^ during the spheroid formation process. (A) Inverted light microscopy (40 x magnification) showing the spheroid formation in agarose micromolds starting from the deposition of 1 · 10^5^ mTECs, (B) Spheroid growth curves from 0 to 108 h, time-point values correspond to the mean and ± SD from three independent replicates. (C) Spheroid cell viability from 0 to 108 h of growth, D) Light microscopy (H&E staining) of histological analysis of spheroid growth from 12 to 48 h, (E) Area of spheroids during its growth from 12 to 48 h. The area was measured and compared through paired student t-test (p-value = *** 0.0007; *** 0.0008; * 0.0368; ns 0.9861 respectively). ns: no significant difference.

### Spheroid dead-cell center and spheroid morphology are impaired in the absence of Aire

In the center of the spheroids, dead cells were observed for both *AIRE*^*WT*^ or *AIRE*^*-/-*^ cells along the growth; however, *AIRE*^*-/-*^ spheroids exhibited an increased rate of dead cells along the growth maximizing at 48 h, as shown in Figure 2A. Quantitative analyzes allowed to determine the relative fluorescence intensity for both live (green-colored) and dead (red-colored) cells (Figure 2B).

**Figure 2A.**
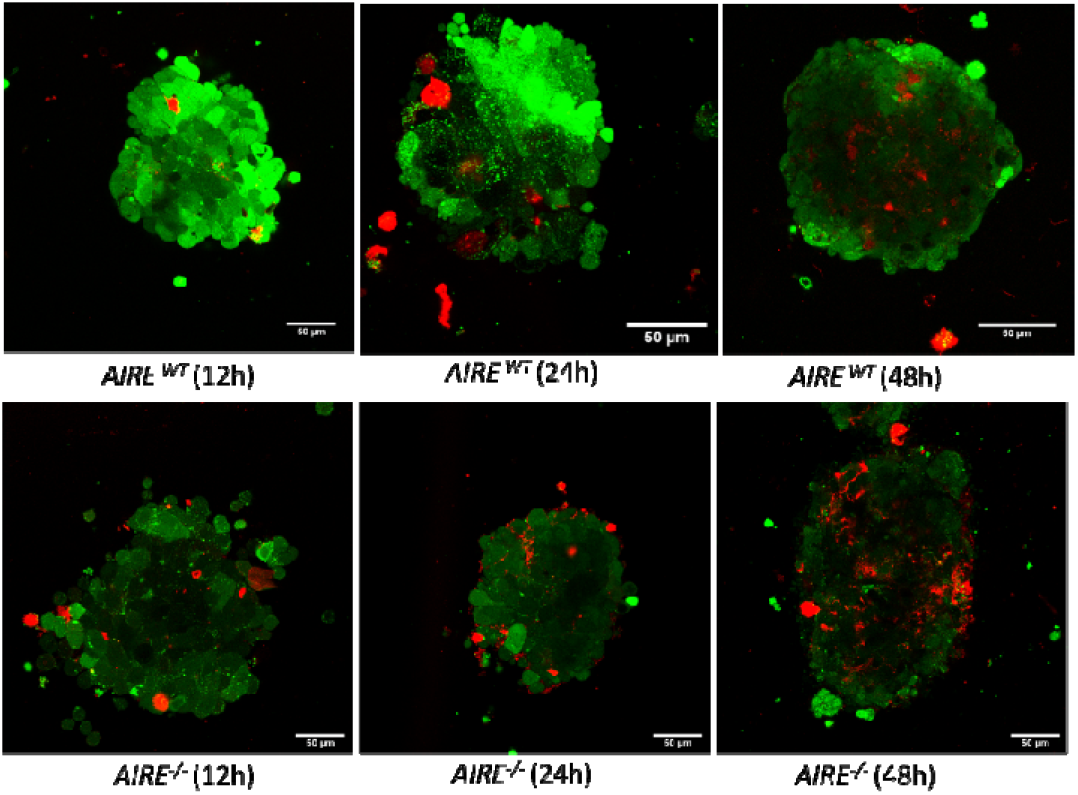

**Figure 2.**
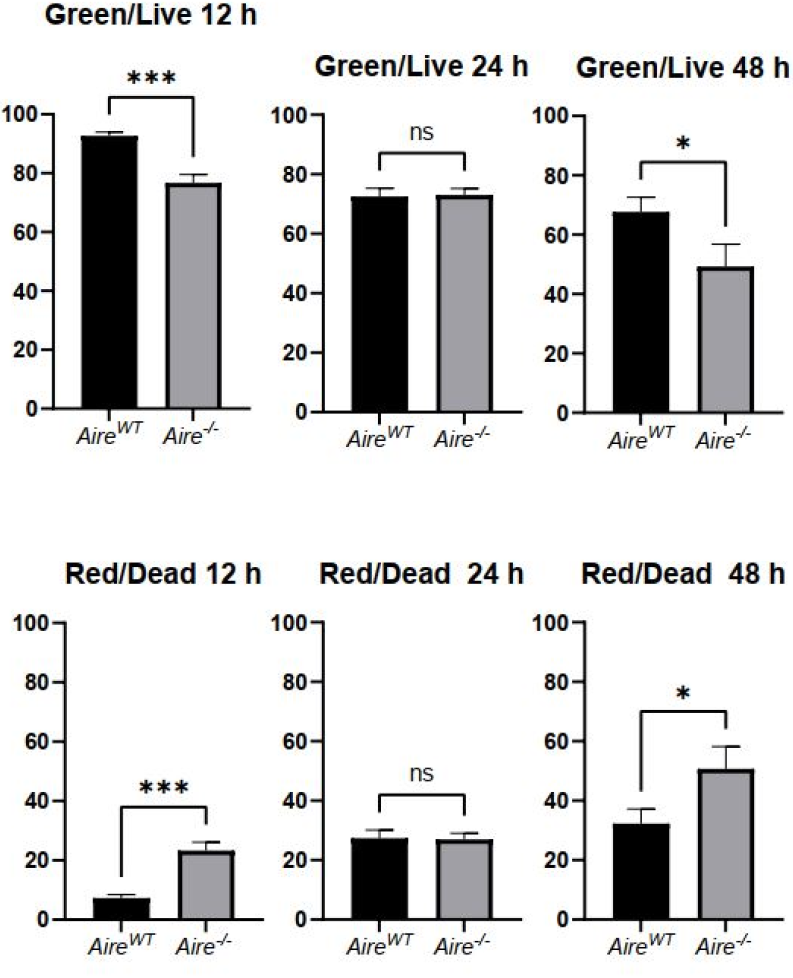
Spheroid confocal fluorescence microscopy. (A) Spheroids were stained with Live/Dead® Viability/Cytotoxicity kit reagents to detect live (green) or dead (red) cells of *Aire*^*WT*^ or A*ire*^*-/-*^ spheroids (B) Quantitative analysis of the fluorescence signal for live or dead cells comparing *Aire*^*WT*^ and *Aire*^*-/-*^ spheroids. Time-point values correspond to mean and ± SD from 20 spheroids analyzed. The MFI of the channels green and red was measured and compared through paired student t-test (p-value = *** 0,0008; ns 0,8097-0,8196; * 0,0237 respectively). ns: no significant difference.

Scanning electron microscopy (SEM) accessed the detailed external surface of the spheroids. Twenty-four-hour *AIRE*^*WT*^ spheroids were more compact exhibiting a well-defined contour, while *AIRE*^*-/-*^ spheroids showed an irregular surface. Throughout SEM, it was possible to observe cells during the adhesion process and visually compare *AIRE*^*WT*^ or *AIRE*^*-/-*^ spheroids (Figure 3).

**Figure 3.**
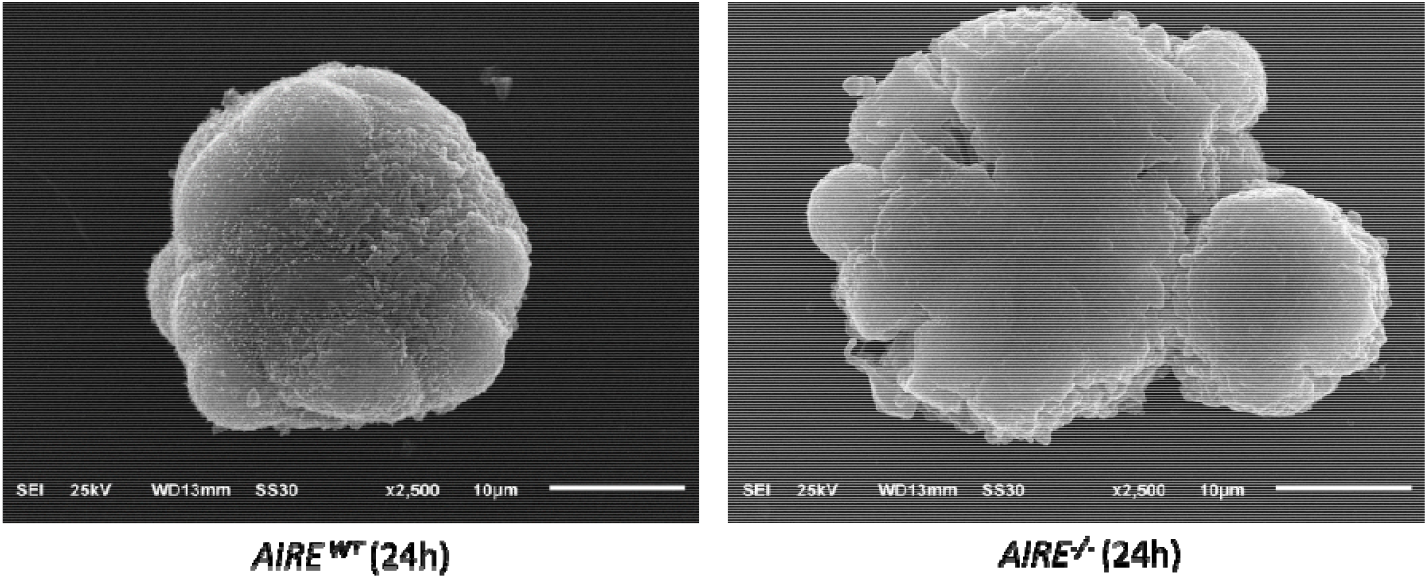
Spheroid scanning electron microscopy. A representative SEM images comparing 24 hours *Aire*^*WT*^ vs. *Aire*^*-/-*^ spheroids. Representative images from 20 spheroids analyzed for each time-point. Scanning electron microscope (Jeol JSM -6610 LV), 5 kV acceleration voltage.

### Phenotypic characterization of spheroid medullary cell surface markers

*AIRE*^*WT*^ spheroids showed: i) the characteristic CD45 Ly51 medullary phenotype, observed in 99.4% of mTECs, ii) 72.9% of the cell population expressed EpCAM and UEA-1, iii) 19.7% of cells were MHC-II, and iv) the CD80 marker was poorly expressed (1.56%) (Figure 4A). In contrast, the *AIRE*^*-/-*^ spheroids showed: i) the characteristic CD45^-^ Ly51^-^ medullary phenotype was observed in 99.3% of mTECs, ii) 14,4% of the cell population expressed EpCAM and UEA-1, iii) 37.7% of cells were CD80, and iv) MHC-II was expressed in 10.2% of the mTECs (Figure 4B).

**Figure 4A.**
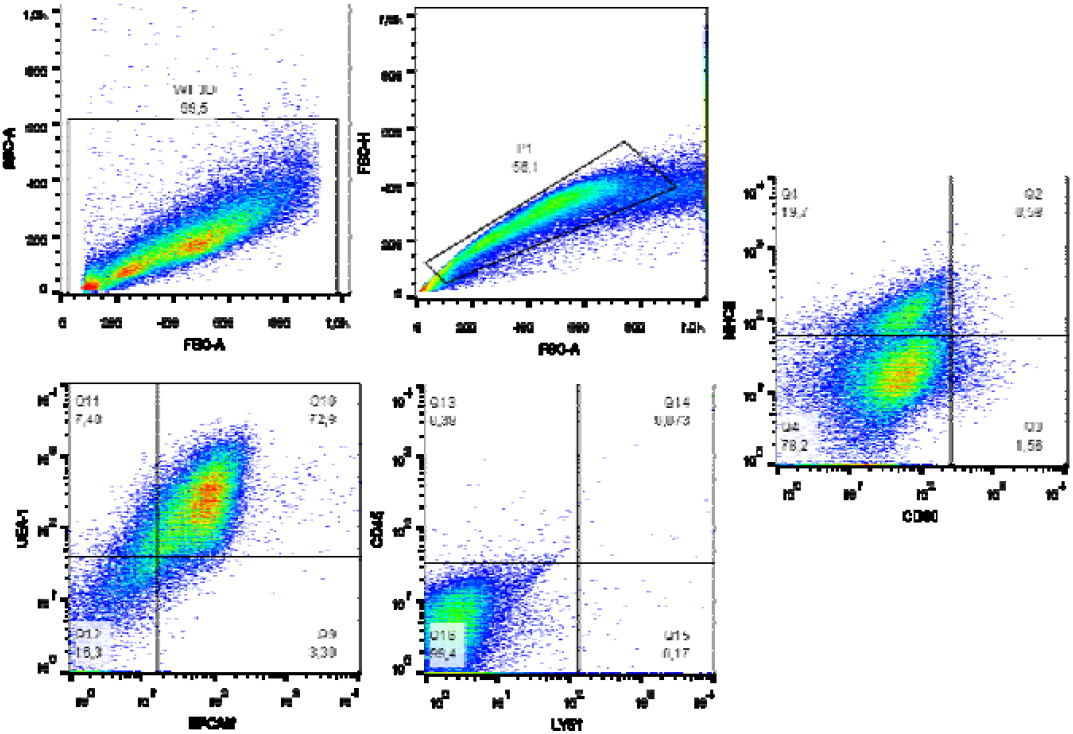

**Figure 4.**
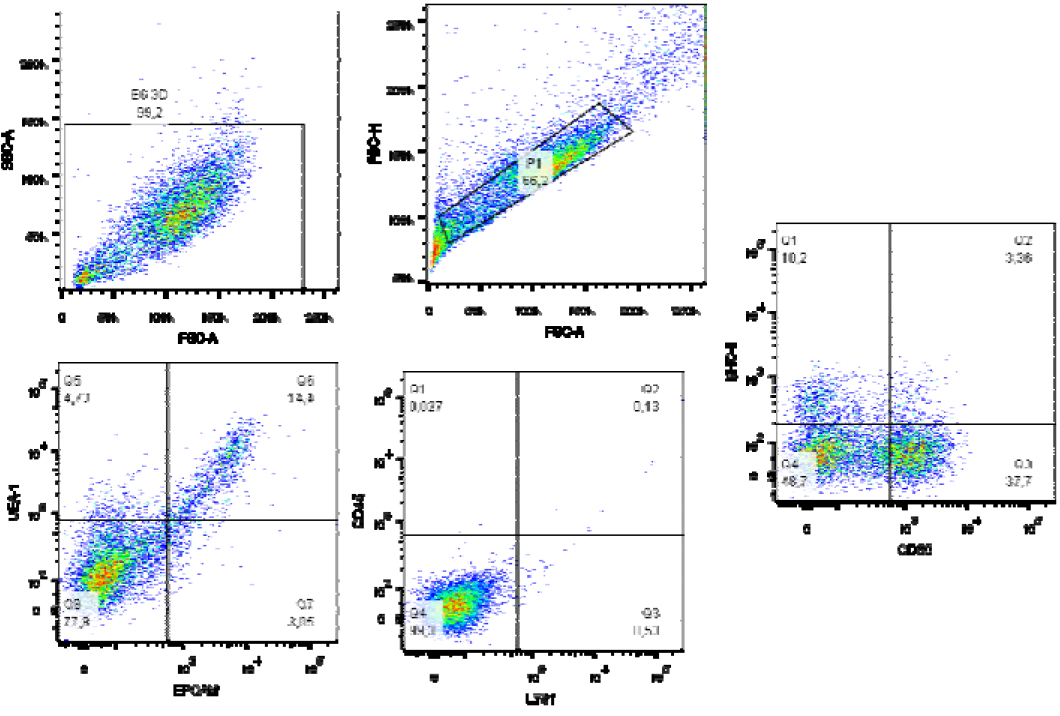
Flow cytometry analysis of phenotype of spheroids *Aire*^*WT*^ or *Aire*^*-/-*^ at 12 h of adhesion. A *AIREWT* spheroids showed the characteristic CD45 Ly51 (99.4%) medullary phenotype, 72,9% of the cell population expressed EpCAM and UEA-1 and 19,7% expressed MHC-II^+^. The CD80 marker was poorly expressed (1,56%). (B) *Aire*^*-/-*^ spheroids showed CD45^-^ Ly51^-^ in 99,3% of the cell population, EpCAM and UEA-1 expressed in 14,4%, CD80 expressed in 37,7% of the cell population and the MHC-II remained poorly expressed (10.2%).

### Spheroid RNA-Seq analysis

Comparative transcriptome analyses were made for the combinations of the study variables at 12 h (*AIRE*^*WT*^ vs. *AIRE*^*-/-*^ spheroids). We identified a set of 1,210 DE mRNAs, of which approximately 83% (1104) correspond to protein-coding mRNAs, and about 17% (106) correspond to non-protein-coding RNAs (Figure 5A). The expression profile of the 1,104 DE mRNAs was determined comparing *AIREWT* vs. *AIRE-/-* spheroids, among which 606 were upregulated and 498 downregulated (Figure 5B). Next, DE mRNAs were hierarchically clustered, and a heat map was constructed to evaluate the individual expression profiles (Figure 5C).

**Figure 5A.**
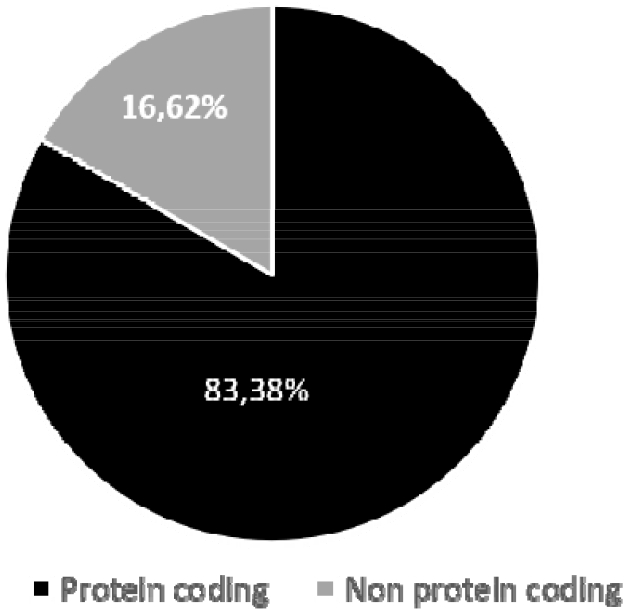

**Figure 5B.**
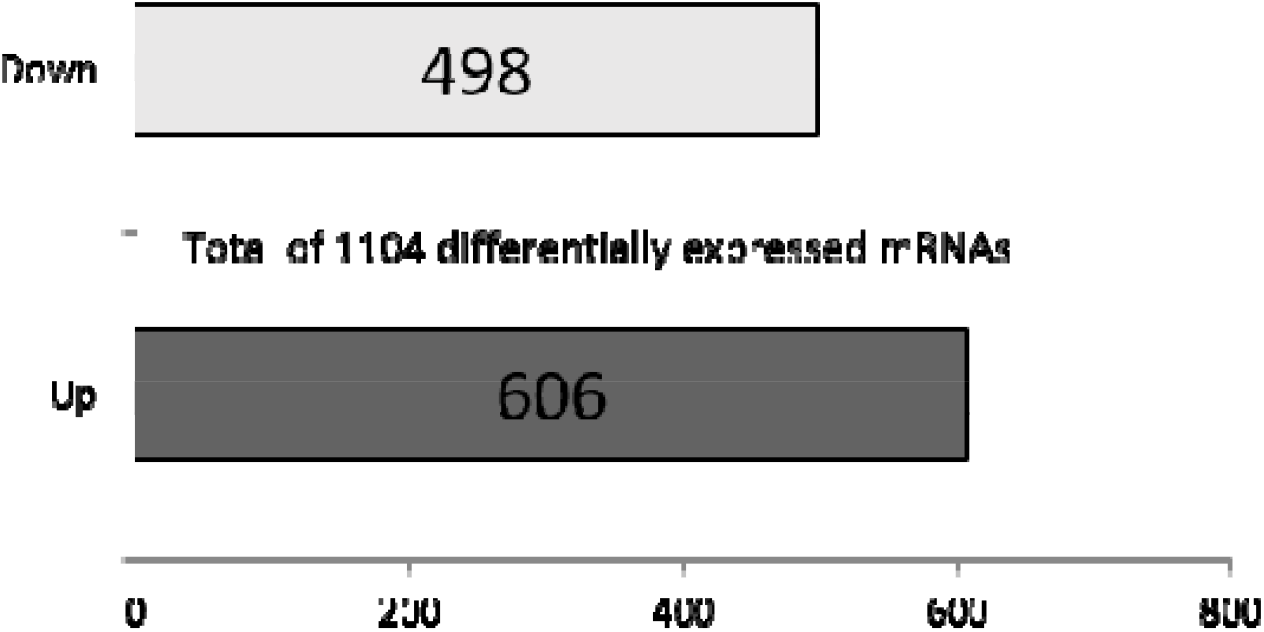

**Figure 5.**
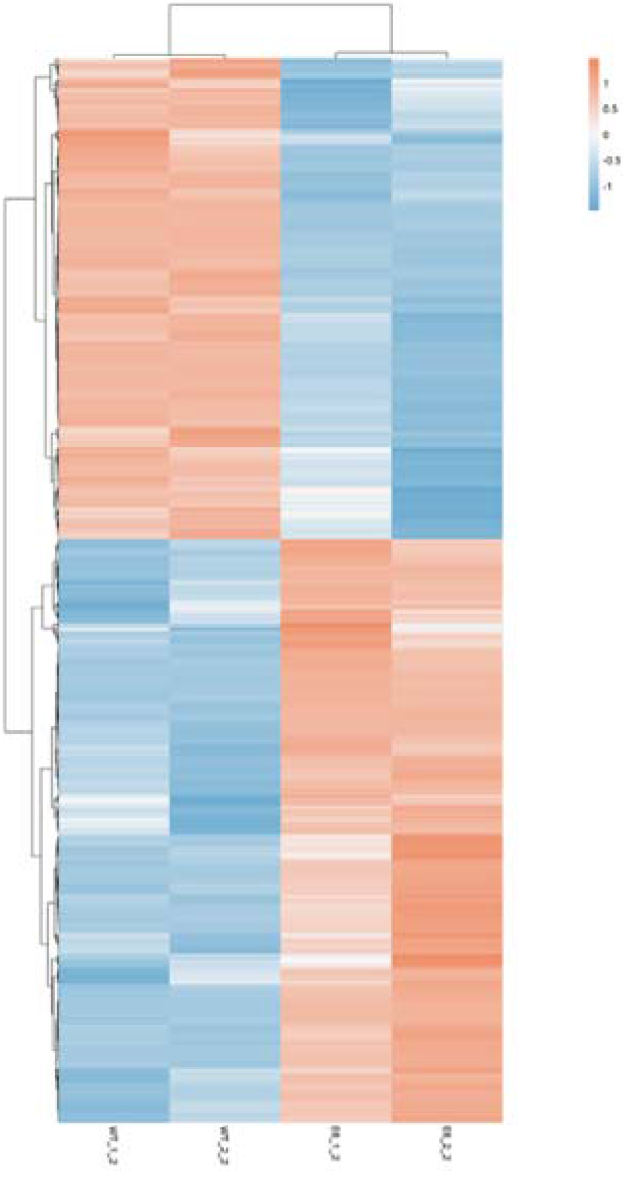
Transcriptome (mRNAs) expression profiles of spheroids. (A) Quantitative analysis of differentially expressed genes grouped to its coding or noncoding function (fold change ≥. 2.0, p-value ≤ 0.05). (B) Number of differentially expressed protein coding genes (fold change ≥. 2.0, p-value ≤ 0.05). (C) Transcriptome (mRNAs) expression profiles of spheroids AIRE^WT^ compared to AIRE^-/-^. The spheroids cells total RNA samples were analyzed through RNA-seq, which allowed identify the differentially expressed mRNAs. The dendrograms and unsupervised heat-maps were obtained using R platform the cluster and tree view algorithm considering 2.0-fold-change and 0.05 false discovery rate. Heat-map legend: red= upregulated, blue= downregulated (Pearson’s correlation metrics).

Figure 6 shows the differences found in the top-ten biological functions of downregulated (Figure 6A) and upregulated mRNAs (Figure 6B).

**Figure 6A.**
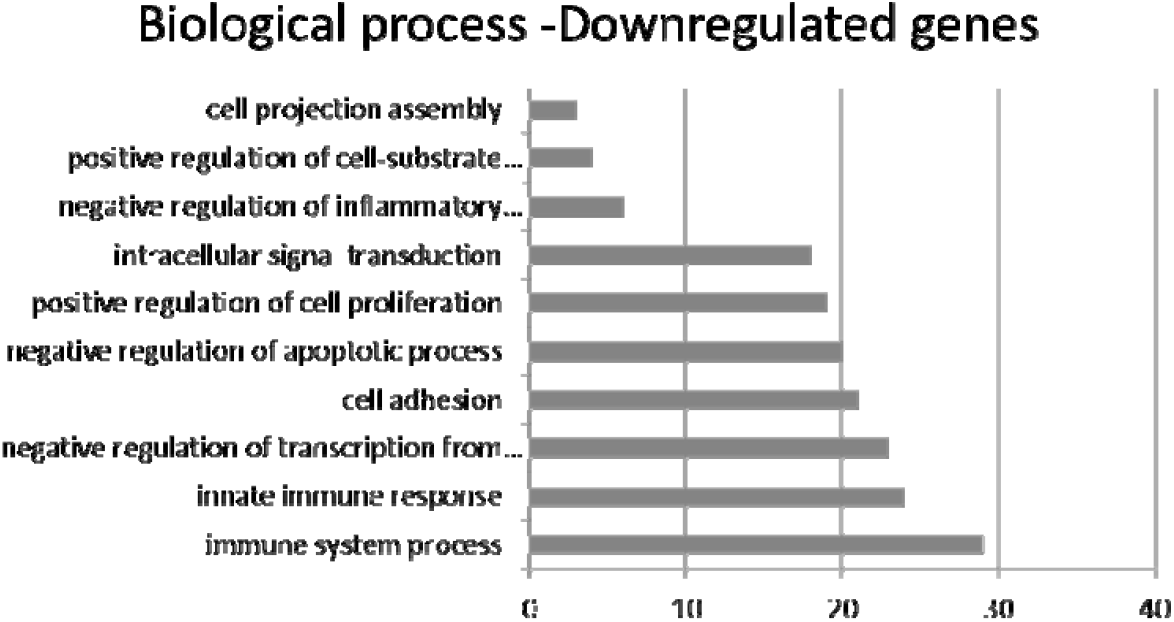

**Figure 6.**
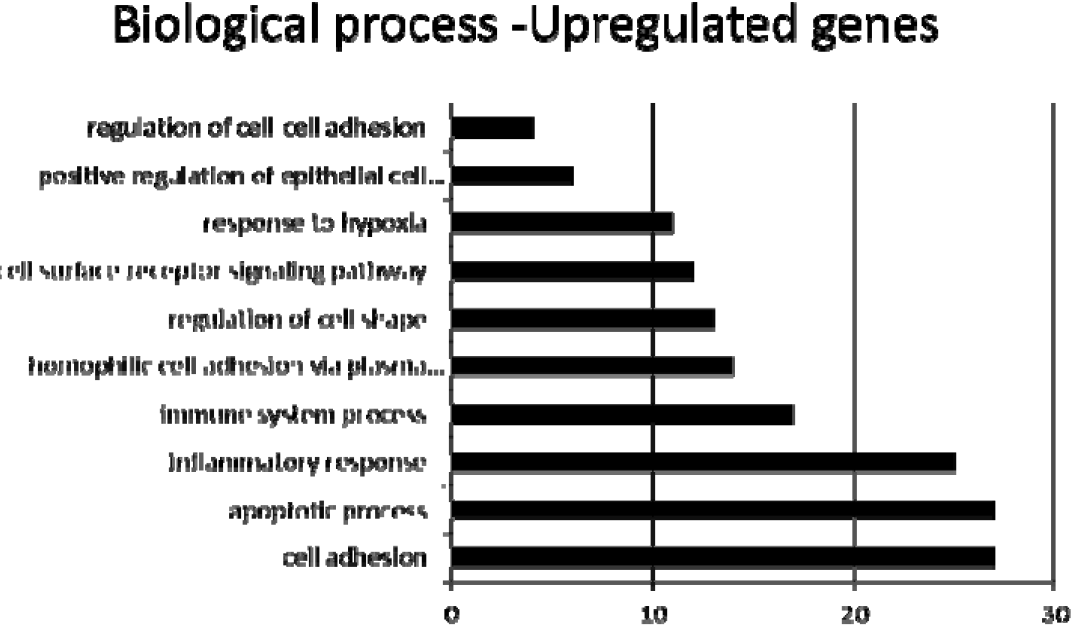
The top 10 biological process involved in spheroid profile comparing Aire WT vs. Aire KO. Functional annotation for modulated down (A) or upregulated (B) mRNAs in spheroids. The functional categories were identified by using DAVID genome database platform according to GO biological processes (BP). The rank is based on the enrichment score, which represents mean *p*-value. Only those mRNA-groups yielding ⍰ 0.05 Benjamini corrected *p*-value and containing at least five mRNAs are considered to be significant (DAVID Bioinformatics Resources Platform 6.8 Database, score <0.05).

Figure 7A show the differentially expressed genes involved in the positive regulation of epithelial cell proliferation. Figure 7B confirms by RT-PCR the differential expression of some top-ten DE mRNAs, including the downregulated *PARVB* that encodes a protein associated with the cell adhesion pathway and the *PLCB2* and *P2RX7* that are involved in the calcium signaling pathway in *AIRE*^*-/-*^ when compared to *AIRE*^*WT*^ spheroids. In contrast, the *VEGFC a*nd *ID1 t*ranscripts, associated with the regulation of epithelial cell proliferation pathway, were upregulated in *AIRE*^*-/-*^ spheroids.

**Figure 7A.**
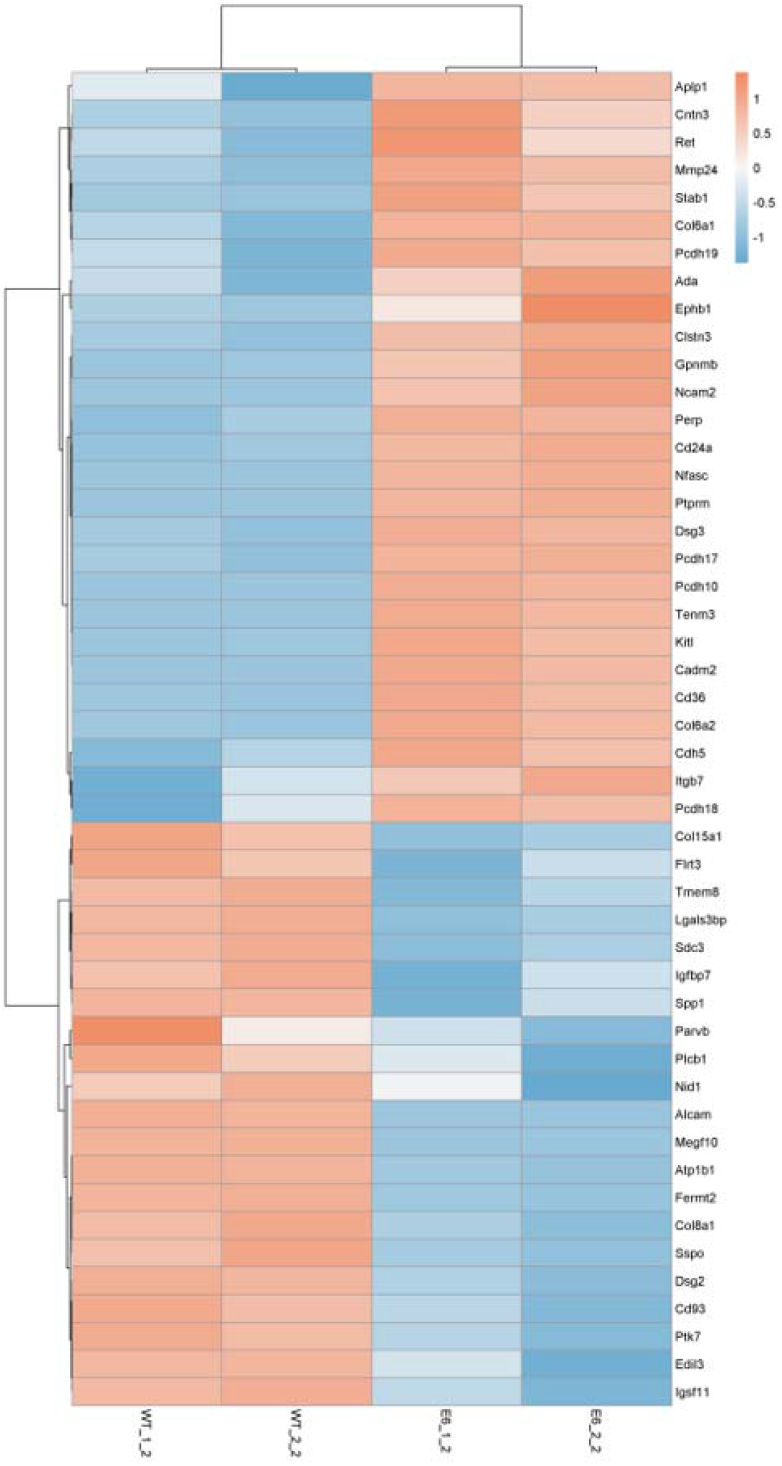

**Figure 7-.**
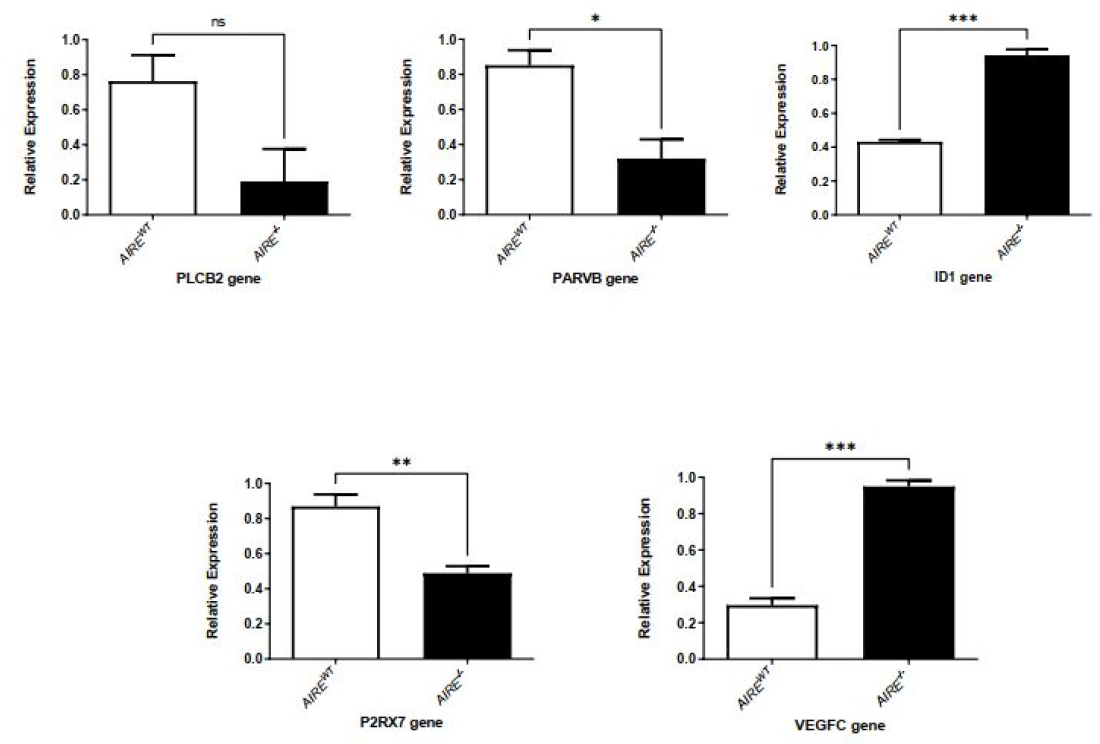
Analysis of RNA sequencing data. (A)-Heat-map showing the large-scale expression profiling of differentially expressed mRNAs involved in cell adhesion pathway. Unsupervised heat-maps and dendrograms were constructed using R platform. Heat-map legend: red= upregulated, blue= downregulated (Pearson’s correlation metrics, fold change ≥ 2.0 and false discovery rate (FDR) <0.05). (B) Evaluation of the relative expression level of the mRNAs PLCB2, PARVB, P2RX7, VEGFC and ID1 by RT-qPCR. Difference between groups was analyzed by Unpaired-t test, comparing data from *AIRE*^*WT*^ vs. *AIRE*^*- /-*^ - p value =PLCB2= ns 0,0738, PARVB= *0,0184, ID1= ***0,0001, P2RX7= ** 0,0089, VEGFC =*** 0,0002.

## Discussion

Considering that: i) the AIRE gene, besides controlling the expression of genes that encode tissue-specific antigens (TSAs) [7, 9, 14], also controls the expression of adhesion molecule genes [17, 18], ii) studies regarding the role of AIRE in the negative selection have primarily focused on the interaction mTEC-thymocyte, and iii) little attention has been devoted to the role of AIRE on the mTEC-mTEC adhesion, in the present study we took advantage of the use of 3D spheroids to evaluate the role of the *AIRE* gene on the mTEC-mTEC adhesion. We showed that the absence of the *AIRE* gene: i) disorganizes the 3D structure of mTEC spheroids, ii) differentially regulates the surface expression of molecules associated with the mTEC phenotype, and iii) differentially modulates the expression of genes encoding adhesion and other molecules.

Several strategies have been developed to study the functional thymic properties, including *ex vivo* reaggregation thymic organ culture (RTOC) [19, 20], 2D mTEC-thymocyte adhesion [17, 18], transwell thymocyte migration [29], 3D fetal thymus organ culture (FTOC) [30], and 3D organotypic culture [23]. Considering that the 3D intrathymic environment is necessary for the development of a functional thymus scaffolding, development of thymic epithelial cells, chemoattraction of pro-T-cells, commitment to the T lineage, and generation of a repertoire of T-cell receptors (TCR) and natural T regulatory cells [31-33], an adequate thymus structure is mandatory to functionality study mTEC-mTEC interactions.

Since the consequences of *AIRE* deletion in the intercellular adhesion remain to be elucidated, we studied the interaction between mTECs when these cells were seeded in a non-adherent substrate such as agarose, where 3D spheroids may be observed at the various stages of development. Once in an agarose substrate, mTECs aggregate and adhere to each other to form spheroids, which may be helpful for the study of mTEC-mTEC adhesion. Then, we evaluated the consequences arising from the absence of Aire, evaluating the: i) mTEC-mTEC physical interaction at regular periods, ii) growth curve of mTEC spheroids, iii) dynamics of 3D mTEC spheroid formation, iv) morphology of mTEC spheroids, v) expression of surface mTEC molecules and vi) gene expression profiles, comparing *AIRE* WT and *AIRE*^*-/-*^ cells.

Our results have shown that Aire interferes in the early process of spheroid formation since, in the absence of Aire, differences in the growth curve and the spheroid morphology were observed. *AIRE*^*WT*^ spheroids were formed at 12 h of culture, while *AIRE*^*-/-*^ spheroids showed a delay in cell-cell adhesion, and spheroid formation occurred only after 20-24 h of culture (Figure 1B/C). Considering that Aire is a pro-apoptotic factor in mature mTECs [34], it is plausible that in addition to controlling mTEC-mTEC adhesion, Aire may also influence the viability, cell cycle, or both during the 3D spheroid formation. Besides, in the model system of this study, it was possible to observe the distinct morphology of the mTEC interactions; i. e., the *AIRE*^*WT*^ forms a typical spheroid at 12 h, whereas the *AIRE*^*-/-*^ initially forms an ellipsoid form instead of a spheroid shape (Figure 1 A/D/E). Although dead cells were observed in the center areas of both spheroid types, the number of these cells progressively increased along time, predominating at 48 h in *AIRE*^*-/-*^ spheroids (Figure 2A-B). Noteworthy, the scanning electron microscopy showed that *AIREWT* spheroids are well-compacted with a well-defined contour. In contrast, *AIRE*^*-/-*^ spheroids exhibited an irregular shape and ill-defined surfaces (Figure 3), indicating that Aire influences the internal and external spheroid structure. Taken in concert, these *in vitro* results corroborate the *in vivo* finding of a disorganized thymic medulla in *AIRE*^*-/-*^ mouse [22] and add information regarding the dynamics of mTEC growth and mTEC-mTEC interactions.

In addition to the spheroid structure, we also evaluate the role of Aire on the mTEC surface markers (Figure 4A/B). Both *AIRE*^*WT*^ or *AIRE*^*-/-*^ spheroids maintain their characteristic CD45^-^ Ly51^-^ medullary profile, i.e., confirming that these cells are not hematopoietic cells (CD45^+^) nor cTECs (Ly51^+^). The absence of AIRE decreased the expression of the double positive EpCAM and UEA-1 cells, which are classical markers for epithelial cells and medullary thymic cells, respectively. Noteworthy, the expression of CD80 was increased in AIRE^-/-^ spheroids (1.57% in *AIRE*^*WT*^ to 37.7% in *AIRE*^*-/-*^) and the expression of MHC-II molecules did not change in both preparations. Three major stages can be observed along mTEC maturation: i) the immature mTEC presents MHC-II^low^, CD80^low^ and Aire^low^, ii) during TRAs, mTECs exhibit their mature stage, presenting MHC-II^high^, CD80^high^ and Aire^high^, and iii) at the post-Aire stage, mTECs are MHC^intermediary^, CD80^intermediary^, accompanied by an increased rate of apoptotic cells [2]. As observed in Figure 4 A/B, the *AIRE*^*WT*^ spheroids presents features of an immature mTEC constitutively exhibiting low expression of AIRE, as previously reported by our group [17]. In contrast, *AIRE*^*-/-*^ spheroids present an intermediate expression of CD80 together with increased rate of dead cells, features that are characteristic of the post-Aire stage. These findings indicate that in the absence of Aire, mTECs present a terminally differentiated mTEC during the first 12 h of 3D spheroid culture, which may affect the interaction with thymocytes.

Considering that Aire controls more than 3,300 genes in mTECs [35], we initially evaluated the mRNA transcript profiles of *AIREWT* or *AIRE-/-*spheroids at 12 hours of mTEC-mTEC adhesion, showing a distinct pattern of gene expression (Figure 5 A-C). Among the DE mRNAs, we identified genes associated with cell adhesion, positive regulation of cell proliferation, apoptotic process, and response to hypoxia (Figure 6 A/B). Among the DE mRNAs related to cell adhesion, we observed a set of transcripts that encode proteins belonging to the cadherins (*PCDHGB7*), collagen (*COL15A1*), integrin (*CIB3*), fibronectin (*FLRT3*), or extracellular matrix protein families (*MMP19)*, which were downregulated in *AIRE-/-*spheroids and putatively involved on the dysregulation of the spheroid formation, as observed in this study. In contrast, *VEGFC* and *ID1* genes involved in the positive regulation of epithelial cell proliferation were upregulated (Figure 7 A/B), a finding that may be associated with the increased number of cells at 12 h of growth (Figure 1B).

Noteworthy, we observed a set of repressed mRNAs associated with the calcium signaling pathway, and among these, the *ADCY3, ADCY1, NOS1, P2RX7, ADRB2, PLCB4, ATP2B4, ATP2A3, P2RX3, CACNA1G, ADRA1B, PLCB1, PLCB2*, and *F2R* transcripts are associated with the intercellular calcium waves. Since intercellular communication used by mTECs is mediated by intercellular calcium waves, requiring functional gap junctions and P2 receptors [36, 37], these genes are potential candidates to be further studied.

Even though operating through different ways, Aire and mTEC-mTEC adhesion culminate into related activities that regulate a cascade of mRNAs that encodes cell adhesion molecules and other relevant molecules that maintain the 3D spheroid formation, opening new perspectives for studies of molecular mechanisms that control the 3D thymic medulla organization and mTEC adhesion and communication.

## Conclusion

While most studies have focused on the mTEC-thymocyte interaction to evaluate the negative selection, this study focused on the role of *AIRE* on the mTEC-mTEC interaction as the primary step to support the adequate thymic medullary structure. Considering the 3D spheroid model, this study reported that the absence of *AIRE* disorganizes the 3D structure of mTEC spheroids, promotes a differentially regulation of mTEC classical surface markers, and modulates genes encoding adhesion and other molecules.

## Acknowledgment

This work was funded by the following Brazilian Research Support Agencies, Fundação de Amparo à Pesquisa do Estado de São Paulo (FAPESP, grants # 17/10780-4 to GAP and EAD, and # 2019/23448-3 to ACM-C), Conselho Nacional de Desenvolvimento Científico e Tecnológico (CNPq, grants # 305787/2017-9 to GAP, and 302060/2019-7 to EAD) and Coordenação de Aperfeiçoamento de Pessoal de Nível Superior (CAPES, Finance Code 001).

We thank Dra. Roberta R. Costa Rosales from the Laboratório Multiusuário de Microscopia Confocal, Ribeirão Preto Medical School, University of São Paulo, Mrs. Maria Dolores and Mr. José Augusto Maulim from the Laboratório Multiusuário de Microscopia Eletrônica, Ribeirão Preto Medical School, and Mrs. Patrícia Viana Bonini Palma and Mrs. Camila Bonaldo, from the Flow Cytometry Laboratory, Ribeirão Preto Blood Center, Ribeirão Preto, SP, Brazil for their technical assistance.

## References

1 Muñoz J.J., Zapata A.G. (2019) Thymus Ontogeny and Development. In: Passos G. (eds) Thymus Transcriptome and Cell Biology. Springer, Cham. https://doi.org/10.1007/978-3-030-12040-5_2

2 Matsumoto M., Rodrigues P.M., Sousa L., Tsuneyama K., Matsumoto M., Alves N.L. (2019) The Ins and Outs of Thymic Epithelial Cell Differentiation and Function. In: Passos G.(eds) Thymus Transcriptome and Cell Biology. Springer, Cham. https://doi.org/10.1007/978-3-030-12040-5_3

3 Abramson, J., & Anderson, G. (2017) .Thymic epithelial cells. Annual Review of Immunology, 35(November 2016), 85–118. https://doi.org/10.1146/annurev-immunol-051116-052320

4 Takaba, H., & Takayanagi, H. (2017) .The Mechanisms of T Cell Selection in the Thymus. Trends in Immunology, 38(11), 805–816. https://doi.org/10.1016/j.it.2017.07.010

5 Cosway, E. J., Lucas, B., James, K. D., Parnell, S. M., Carvalho-Gaspar, M., White, A. J., Tumanov, A. V., Jenkinson, W. E., & Anderson, G. (2017) .Redefining thymus medulla specialization for central tolerance. Journal of Experimental Medicine, 214(11), 3183–3195. https://doi.org/10.1084/jem.20171000

6 Yoganathan K., Chen E.L.Y., Singh J., Zúñiga-Pflücker J.C. (2019) T-Cell Development: From T-Lineage Specification to Intrathymic Maturation. In: Passos G. (eds) Thymus Transcriptome and Cell Biology. Springer, Cham. https://doi.org/10.1007/978-3-030-12040-5_4

7 Derbinski, J., Pinto, S., Rösch, S., Hexel, K., & Kyewski, B. (2008) .Promiscuous gene expression patterns in single medullary thymic epithelial cells argue for a stochastic mechanism. Proceedings of the National Academy of Sciences of the United States of America, 105(2), 657–662. https://doi.org/10.1073/pnas.0707486105

8 Abramson, J., Giraud, M., Benoist, C., & Mathis, D. (2010) .Aire’s Partners in the Molecular Control of Immunological Tolerance. Cell, 140(1), 123–135. https://doi.org/10.1016/j.cell.2009.12.030

9 Passos, G. A., Speck-Hernandez, C. A., Assis, A. F., & Mendes-da-Cruz, D. A. (2018) .Update on Aire and thymic negative selection. Immunology, 153(1), 10–20. https://doi.org/10.1111/imm.12831

10 Passos, G. A., Genari, A. B., Assis, A. F., Monteleone-, A. C., Donadi, E. A., Oliveira, E. H., Duarte, M. J., Machado, M. V, Tanaka, P. P., & Mascarenhas, R. (2019) .The Thymus as a Mirror of the Body’s Gene Expression. In: Passos G. (eds) Thymus Transcriptome and Cell Biology. Springer, Cham. https://doi.org/10.1007/978-3-030-12040-5_9

11 St-Pierre, C., Brochu, S., Vanegas, J. R., Dumont-Lagacé, M., Lemieux, S., & Perreault, C. (2013) .Transcriptome sequencing of neonatal thymic epithelial cells. Scientific Reports, 3. https://doi.org/10.1038/srep01860

12 Sansom, S. N., Shikama-Dorn, N., Zhanybekova, S., Nusspaumer, G., Macaulay, I. C., Deadman, M. E., Heger, A., Ponting, C. P., & Holländer, G. A. (2014) .Population and single-cell genomics reveal the Aire dependency, relief from Polycomb silencing, and distribution of self-antigen expression in thymic epithelia. Genome Research, 24(12), 1918–1931. https://doi.org/10.1101/gr.171645.113

13 Abramson, J., & Goldfarb, Y. (2016) .AIRE: From promiscuous molecular partnerships to promiscuous gene expression. European Journal of Immunology, 46(1), 22–33. https://doi.org/10.1002/eji.201545792

14 Irla M. (2019) Thymic Crosstalk: An Overview of the Complex Cellular Interactions That Control the Establishment of T-Cell Tolerance. In: Passos G. (eds) Thymus Transcriptome and Cell Biology. Springer, Cham. https://doi.org/10.1007/978-3-030-12040-5_6

15 Giraud, M., Yoshid, H., Abramson, J., Rahl, P. B., Young, R. A., Mathis, D., & Benoist, C. (2012) .Aire unleashes stalled RNA polymerase to induce ectopic gene expression in thymic epithelial cells. Proceedings of the National Academy of Sciences of the United States of America, 109(2), 535–540. https://doi.org/10.1073/pnas.1119351109

16 Takaba, H., Morishita, Y., Tomofuji, Y., Danks, L., Nitta, T., Komatsu, N., Kodama, T., & Takayanagi, H. (2015) .Fezf2 Orchestrates a Thymic Program of Self-Antigen Expression for Immune Tolerance. Cell, 163(4), 975–987. https://doi.org/10.1016/j.cell.2015.10.013

17 Pezzi, N., Assis, A. F., Cotrim-Sousa, L. C., Lopes, G. S., Mosella, M. S., Lima, D. S., Bombonato-Prado, K. F., & Passos, G. A. (2016) .Aire knockdown in medullary thymic epithelial cells affects Aire protein, deregulates cell adhesion genes and decreases thymocyte interaction. Molecular Immunology, 77, 157–173. https://doi.org/10.1016/j.molimm.2016.08.003

18 Speck-Hernandez, C. A., Assis, A. F., Felicio, R. F., Cotrim-Sousa, L., Pezzi, N., Lopes, G. S., Bombonato-Prado, K. F., Giuliatti, S., & Passos, G. A. (2018) .Aire disruption influences the medullary thymic epithelial cell transcriptome and interaction with thymocytes. Frontiers in Immunology, 9(MAY). https://doi.org/10.3389/fimmu.2018.00964

19 White, A., Jenkinson, E., & Anderson, G. (2008) .Reaggregate thymus cultures. Journal of Visualized Experiments, 18. https://doi.org/10.3791/905

20 Tajima, Asako, et al. “Promoting 3-D Aggregation of FACS Purified Thymic Epithelial Cells with EAK 16-II/EAKIIH6 Self-assembling Hydrogel.” Journal of visualized experiments: JoVE 112 (2016).

21 Mendes-Da-Cruz, D. A., Stimamiglio, M. A., Muñoz, J. J., Alfaro, D., Terra-Granado, E., Garcia-Ceca, J., Alonso-Colmenar, L. M., Savino, W., & Zapata, A. G. (2012) .Developing T-cell migration: Role of semaphorins and ephrins. FASEB Journal, 26(11), 4390–4399. https://doi.org/10.1096/fj.11-202952

22 Ramsey, C., Winqvist, O., Puhakka, L., Halonen, M., Moro, A., Kämpe, O., Eskelin, P., Pelto-Huikko, M., & Peltonen, L. (2002) .Aire deficient mice develop multiple features of APECED phenotype and show altered immune response. Human Molecular Genetics, 11(4), 397–409. https://doi.org/10.1093/hmg/11.4.397

23 Pinto, S., Stark, H. J., Martin, I., Boukamp, P., & Kyewski, B. (2015) .3D Organotypic Co-Culture Model Supporting Medullary Thymic Epithelial Cell Proliferation, Differentiation and Promiscuous Gene Expression. Journal of Visualized Experiments, 2015(101), 1–7. https://doi.org/10.3791/52614

24 Ucar, A., Ucar, O., Klug, P., Matt, S., Brunk, F., Hofmann, T. G., & Kyewski, B. (2014) .Adult thymus contains foxN1-epithelial stem cells that are bipotent for medullary and cortical thymic epithelial lineages. Immunity, 41(2), 257–269. https://doi.org/10.1016/j.immuni.2014.07.005

25 Sheridan, J. M., Keown, A., Policheni, A., Roesley, S. N. A., Rivlin, N., Kadouri, N., Ritchie, M. E., Jain, R., Abramson, J., Heng, T. S. P., & Gray, D. H. D. (2017) .Thymospheres Are Formed by Mesenchymal Cells with the Potential to Generate Adipocytes, but Not Epithelial Cells. Cell Reports, 21(4), 934–942. https://doi.org/10.1016/j.celrep.2017.09.090

26 Silva, C. S., Pinto, R. D., Amorim, S., Pires, R. A., Correia-Neves, M., Reis, R. L., Alves, N. L., Martins, A., & Neves, N. M. (2020) .Fibronectin-Functionalized Fibrous Meshes as a Substrate to Support Cultures of Thymic Epithelial Cells. Biomacromolecules, 21(12), 4771–4780. https://doi.org/10.1021/acs.biomac.0c00933

27 Hirokawa K, Utsuyama M, Moriizumi E, Handa S. Analysis of the thymic microenvironment by monoclonal antibodies with special reference to thymic nurse cells. Thymus. 1986 ;8(6):349–360. PMID: 3101237.

28 Mizuochi, T., Kasai M, Kokuho T, Kakiuchi T, and Hirokawa K (1992). Medullary but not cortical thymic epithelial cells present soluble antigens to helper T cells. Journal of Experimental Medicine, 175, 1601–1605. https://doi.org/10.1084/jem.175.6.1601

29 Pérez, A. R., Berbert, L. R., Lepletier, A., Revelli, S., Bottasso, O., Silva-Barbosa, S. D., & Savino, W. (2012) .TNF-α is involved in the abnormal thymocyte migration during experimental trypanosoma cruzi infection and favors the export of immature cells. PLoS ONE, 7(3). https://doi.org/10.1371/journal.pone.0034360

30 Anderson, G., & Jenkinson, E. J. (2007) .Fetal Thymus Organ Culture. Cold Spring Harbor Protocols, 2007(8), ppdb.prot4808-pdb.prot4808. https://doi.org/10.1101/pdb.prot4808

31 Kyewski, B., & Klein, L. (2006) .A central role for central tolerance. Annual Review of Immunology, 24, 571–606. https://doi.org/10.1146/annurev.immunol.23.021704.115601

32 Hogquist, K. A., Baldwin, T. A., & Jameson, S. C. (2005) .Central tolerance: Learning self-control in the thymus. Nature Reviews Immunology, 5(10), 772–782. https://doi.org/10.1038/nri1707

33 Brunk, F., Michel, C., Holland-Letz, T., Slynko, A., Kopp-Schneider, A., Kyewski, B., & Pinto, S. (2017) .Dissecting and modeling the emergent murine TEC compartment during ontogeny. European Journal of Immunology, 47(7), 1153–1159. https://doi.org/10.1002/eji.201747006

34 Sun, L., Li, H., Luo, H., & Zhao, Y. (2014) .Thymic epithelial cell development and its dysfunction in human diseases. BioMed Research International, 2014. https://doi.org/10.1155/2014/206929

35 St-Pierre, C., Trofimov, A., Brochu, S., Lemieux, S., & Perreault, C. (2015) .Differential Features of AIRE-Induced and AIRE-Independent Promiscuous Gene Expression in Thymic Epithelial Cells. The Journal of Immunology, 195(2), 498–506. https://doi.org/10.4049/jimmunol.1500558

36 Da, R., Bisaggio, C., Nihei, O. K., Persechini, P. M., Fundação, W. S., & Cruz, O. (2017) .Characterization of P2 receptors in thymic epithelial cells. https://www.researchgate.net/publication/12042294

37 Nihei, O. K., Campos de Carvalho, A.C., Spray, D. C., Savino, W., & Alves, L. A. (2003) .A novel form of cellular communication among thymic epithelial cells: Intercellular calcium wave propagation. American Journal of Physiology - Cell Physiology, 285(5 54-5), 1304–1313. https://doi.org/10.1152/ajpcell.00568.2002

